# Applying expression profile similarity for discovery of patient-specific functional mutations

**DOI:** 10.1101/172015

**Authors:** Guofeng Meng

## Abstract

The progress of cancer genome sequencing projects yields unprecedented information of mutations for numerous patients. However, the complexity of mutation profiles of patients hinders the further understanding of mechanisms of oncogenesis. One basic question is how to uncover mutations with functional impacts. In this work, we introduce a computational method to predict functional somatic mutations for each of patient by integrating mutation recurrence with similarity of expression profiles of patients. With this method, the functional mutations are determined by checking the mutation enrichment among a group of patients with similar expression profiles. We applied this method to three cancer types and identified the functional mutations. Comparison of the predictions for three cancer types suggested that most of the functional mutations were cancer-type-specific with one exception to p53. By checking prediction results, we found that our method effectively filtered non-functional mutations resulting from large protein sizes. In addition, this methods can also perform functional annotation to each patient to describe their association with signalling pathways or biological processes. In breast cancer, we predicted “cell adhesion” and other mutated gene associated terms to be significantly enriched among patients.

## 1 Introduction

Cancer arises from accumulations of somatic mutations and other genetic alterations, which leads to abnormal cell proliferation [1]. With the progress of cancer genome sequencing projects, such as The Cancer Genome Atlas (TCGA) project, mutation information for more and more patients are becoming publicly available [2, 3, 4], which provides the foundation to uncover mechanisms of oncogenesis. However, cancer patients usually carry an average of 33 to 66 mutations in the protein coding regions and those mutations are supposed to take unequal roles in oncogenesis [5, 6]. It remains a great challenge to distinguish the functional mutations that give cells with growth advantages [7], from the ones with non-crucial roles to oncogenesis.

Many computational tools have been developed to predict the functional mutations. One popular strategy is to find recurrently mutated genes. Genes with higher mutation frequency are supposed to take a more essential role in oncogenesis [6, 8, 9]. To improve the accuracy, most computational tools also consider background mutation rates and protein sizes in discovery of recurrent mutations [10, 11, 12]. Another strategy is mutual exclusivity, which is based on an assumption that if one gene is mutated in a patient, other members acting in the same signalling pathway are less likely to be mutated [13]. Using mutual exclusivity, members of signalling pathways are investigated for their coverage across a large number of patients while not co-mutated in the same patients [14, 15, 16, 17]. Besides of functional annotations, expression information is also used in driver mutation discovery. One popular application is to find functional copy number variations (CNVs) by identifying the differentially expressed genes located in the CNV regions [18]. For somatic mutations, correlation of gene expression profiles is also used to identify the functional mutations based on the assumption that somatic mutations will result in altered expression of their downstream targets [19].

Somatic mutations in coding regions are usually supposed to involve oncogenesis by affecting the activities of proteins associated with cell proliferation [20]. However, somatic mutations at different sites have different effects on protein activities. One solution is to study the stability of protein structures after mutations. Amino acid changes in the protein sequences can either stabilize, destabilize or have no effect on protein structures. Methods based on this strategy calculate the protein folding free energy, which will be used to evaluate the protein structural stability [21, 22, 23]. However, those methods still face the problems of establishing the direct connections between protein structures and activities.

In this work, we introduce a novel method to recover functional somatic mutations for each of the studied patients by integrating mutation recurrence and expression profile similarity among patients. This method is based on two assumptions: (1) functional mutations will lead to altered expression of genes which reflects consequences of somatic gene mutations; (2) patients with similar expression profiles are more likely to carry functional mutations to the same genes. For each patient, we can find a group of patients that have similar expression profiles to him/her and those patients are called neighboring patients. Mutations of the studied patient are evaluated for their enrichment among all the neighboring patients. Mutations with enough enrichment are predicted to be functional mutations. The functional mutations of all patients can be recovered by repeating this process. This method also performs function annotation analysis to mutated genes in neighboring patients so that each patient can be assigned with some functional terms to indicate his/her association with signalling pathways or biological processes. As applications, we applied this method to three cancer data sets and identified the functional mutations for three type of cancers respectively.

## 2 Materials and Methods

### 2.1 Dataset and their pre-processing

In this work, we focus on prediction of functional somatic mutations, including missense mutation, nonsense mutation and frame-shift. All the mutation and expression data of cancer patients used in this work are downloaded from TCGA project (by May 2013) at level two, with which the expression data have been normalized within samples and that the somatic mutations have also been annotated in exons by the pilot of TCGA project. We exclude those patients with only mutation or expression data for next-step analysis.

### 2.2 Expression biomarkers

To describe the expression profiles of patients, we firstly determine a group of genes or probes as expression biomarkers by selecting the top 2000 genes or probes with the most expression variances among all the patients. These biomarkers are supposed to better reflect the expression consequences of gene mutations. The expression profile of one patient is described as a vector with the expression values of biomarkers as the elements.

### 2.3 Neighboring patients: patients with similar expression profiles

The similarity of expression profiles of the patients was measured by using Pearson’s correlation *r*. Conversely, the distances between patients could be described by 1 – *r*. For each patient, we could find his/her neighboring patients by selecting *n* patients with most significant positive correlation values at a minimum *r* cutoff ( e.g. *r* > 0.6), where *n* ranged from 20 to 50.

## 3 Results

### 3.1 Association of mutation and expression profiles

As an exemplary illustration, we checked the association between somatic mutations and cancer patients with similar expression profiles by investigating 516 breast invasive cancer patients, for which both expression and somatic mutation information are available from the TCGA project. We selected 2000 probes with most expression variances as the biomarkers and performed hierarchical clustering to the patients. Then we checked the distribution of mutated genes on the hierarchical tree. One example is CDH1 gene, which is mutated in 35 out of 516 patients. As shown in Figure 1, patients with CDH1 gene mutation were not randomly distributed but preferentially clustered together. To quantify this, we classified the patients into six groups based on the structure of hierarchical clustering tree. Among groups, the frequency of CHD1 mutations varied greatly. In Cluster IV, we observed 22 out of 120 patients to carry CDH1 mutations, which was also the most enriched group. Fisher’s exact test indicated the significance of this enrichment to be *p* < 4.4*e* – 8. In cluster VI, there were 6 out of 70 patients to carry CDH1 mutation without statistical significance. For cluster I and V, there was even no patient with CDH1 mutation. We also checked the mutation distribution of other genes and observed non-random distribution in the expression subclusters in nearly all the test cases.

**Figure 1:**
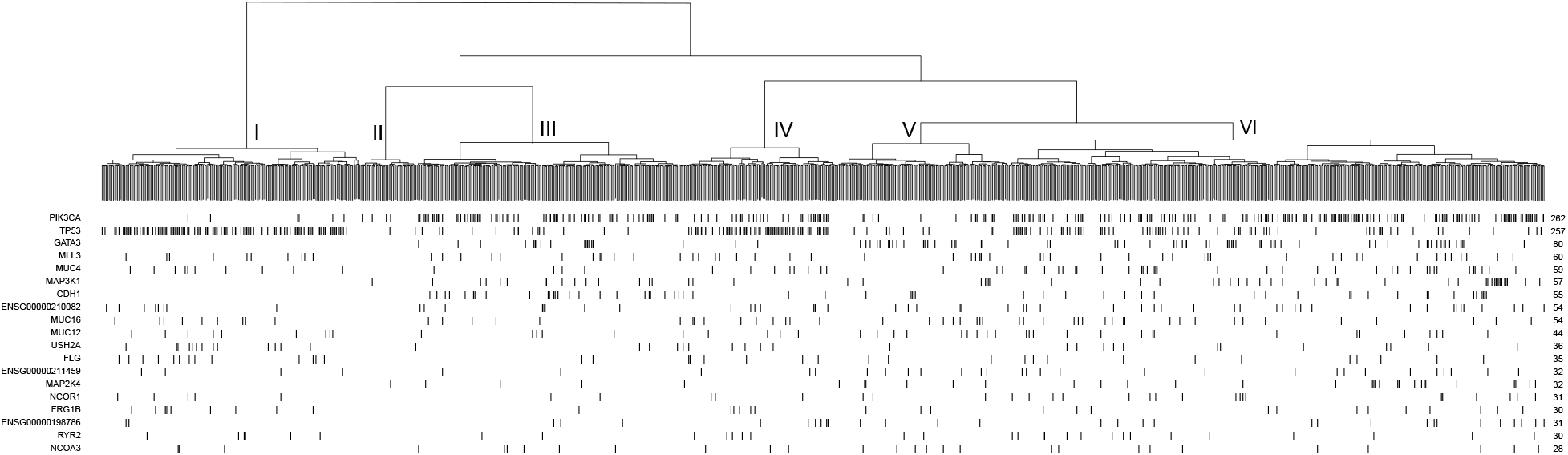
Hierachical clustering and the mutation distribution of gene

These results suggest that patients with a similar set of mutated genes are more likely to have similar expression profiles, which can be explained by the fact that gene mutations would directly or indirectly affect transcription of their downstream targets [19, 24]. Based on this observation, we propose that genes affected by functional mutations can be identified by checking their enrichment in patients with similar expression profiles.

### 3.2 Pipeline to find functional mutations

In this work, we propose a computational method to infer the functional mutations for each studied patient. This method is based on two assumptions that, (1) genes carrying functional mutations directly or indirectly lead to altered expression of their downstream target genes; (2) patients with the same functional mutations will share similar expression profiles. Thus, for each mutated gene from one patient, we can check its mutation enrichment in the neighboring patients. The impact of gene mutation to studied patients can be described by a *p*-value based on statistical frameworks.

The whole process is described in Figure 2. This method requires two matrices as input: one expression matrix and one binary mutation matrix. As described in Section Methods, the expression matrix are explored to determine the neighbor patients by checking its expression similarity. For patient *P*_1_ with a mutation to gene *g_i_*, his/her neighboring patients are selected by choosing *n* patients (including *P*_1_) with the most similar expression profiles with *P*_1_, where *n* ranged from 5 to 30 (a). Then, the patient number with mutation to gene *g_i_* is counted in all the neighbors of *P*_1_ (b). Based on randomly simulation, in which the neighbor of patients are randomly selected, the enrichment of mutations to *g_i_* among neighbor patients is evaluated by Fisher’s exactly test. A *p*-value is assigned to studied patient to describe its mutation enrichment to *g_i_* (c), which also indicates its functional importance. If the *p*-value is significant enough (e.g. 0.01), mutation to gene *g_i_* in patient *P*_1_ will be treated as functional mutation of only patient *P*_1_. The same process can be repeated for other mutated genes and patients one by one (f).

**Figure 2:**
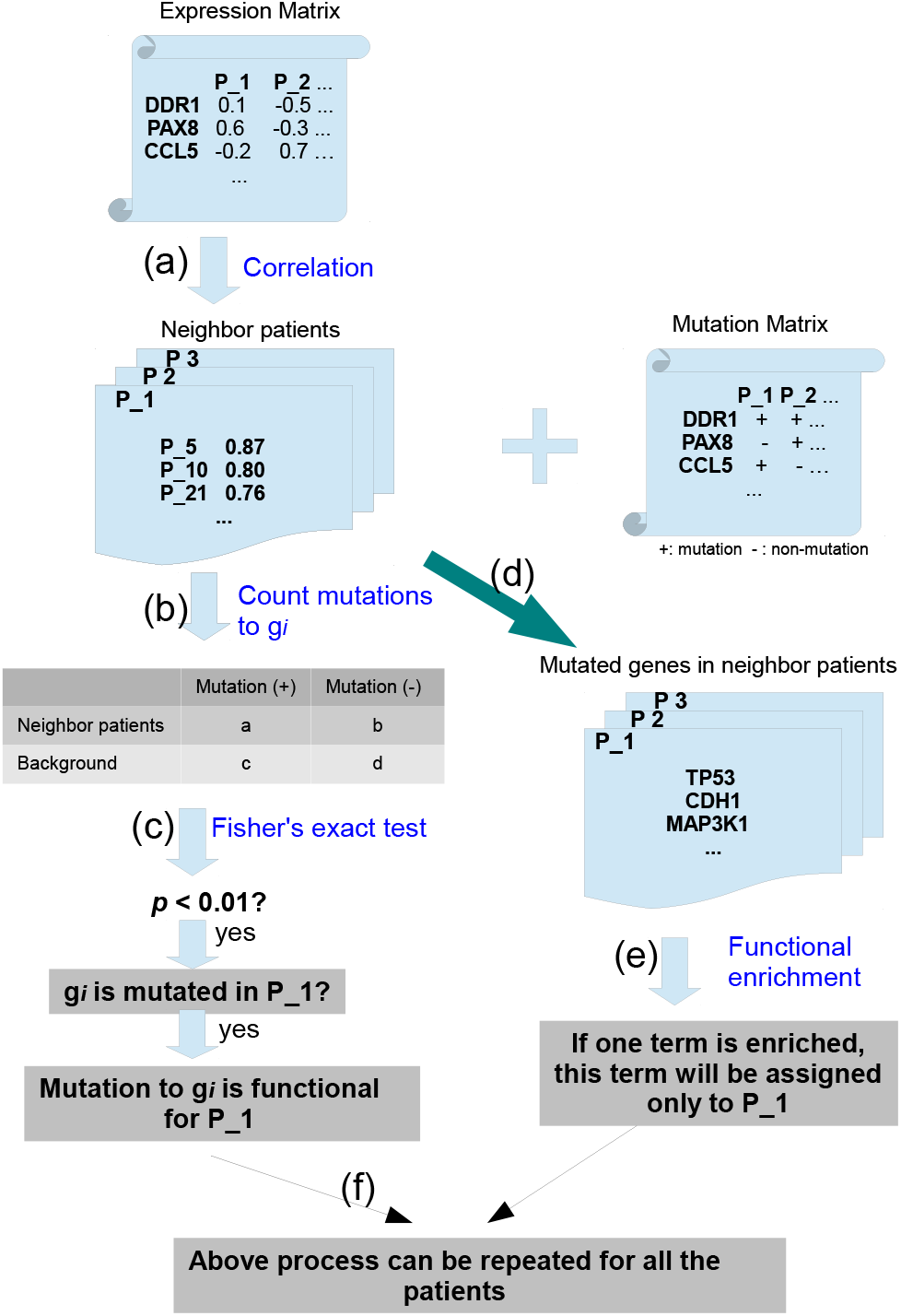
Pipeline to predict functional mutations and terms (see main text for detailed description).

With our method, we also check the functional annotation for each studied patient by enrichment analysis to all the mutated genes in neighboring patients. The functional annotation is described by Gene ontology (GO) or KEGG terms. In this step, all the mutated genes from patient *P*_1_ and his/her neighboring patients (d) are checked for enriched functional terms by comparing to the background occurrences with Fisher’s exact test (e). If a term is significantly enriched (e.g. *p* < 0.01), it will be assigned to patient *P*_1_ as functional annotation. This term indicates the functional consequences of gene mutations in patient *P*_1_. For the other patients, the same process can be repeated to find their associated functional terms (f).

### 3.3 Application to breast cancer

As an application, we applied our method to 516 beast cancer patients from the TCGA project. The functional mutations were chosen at a cutoff of *p* < 0.01. In Figure 1 (1), we show 19 genes with the most mutation recurrences in breast cancer. For all the genes, only part of their mutations are predicted to be functional. Among them, PIK3CA and TP53 were especially recurrently mutated based on our prediction. PIK3CA gene was observed to be mutated in 175 breast cancer patients and 123 of them were confirmed to be functional (about 70.3%), which made PIK3CA to be the gene with the most confirmed mutations, even though it was not the one with the most somatic mutations in breast cancer. Another gene was TP53 with functional mutations in 107 patients, about 56.9% of observed somatic mutations. For other genes, relatively fewer functional mutations were observed compared to PIK3CA and TP53. However, many of them have been widely reported for their critical roles in breast cancer, such as MAP3K1 [25], CDH1 [26], GATA3 [27], which indicated their important roles but in fewer patients. Overall, we observed 408 of 516 breast cancer patients to carry at least one of these 10 predicted driver mutations, covering about 79% of all the patients.

**Table 1:** Top 10 of functional mutations in breast cancer

| Mutated gene | No. somatic mutation | No. functional mutation | Percentage |
| --- | --- | --- | --- |
| PIK3CA | 175 | 123 | 70.3% |
| TP53 | 188 | 107 | 56.9% |
| MAP3K1 | 40 | 30 | 75.0% |
| CDH1 | 35 | 29 | 82.6% |
| GATA3 | 56 | 19 | 33.9% |
| RUNX1 | 19 | 13 | 68.4% |
| CTCF | 15 | 10 | 66.7% |
| CACNA1B | 14 | 9 | 64.2% |
| DNAH17 | 14 | 8 | 57.1% |
| MAP2K4 | 21 | 8 | 38.1% |

Next, we tested whether our predicted mutations were more likely to be functional. We checked genes with recurrent somatic mutations while less often predicted to be functional. One example was the TTN gene with somatic mutation in 90 patients, which ranked as the gene with the third most recurrent mutations. However, we only predicted three of them to be functional. TTN protein is a component of vertebrate striated muscle [28]. By searching literature, we did not find any reports to support TTN to take critical role in cancer. By checking its protein sequence, we found that it is very long with a length of 27,000 to 33,000 amino acids [29], indicating a recurrence bias of its mutations. This guess was also confirmed by the analysis with MutSig, which considered the protein size and background mutation rates into evaluation [10]. Similar results are observed with with other genes, such as MUC6 with mutations in 57 patients while predicted to be functional in only two patients. Even though our pipeline did not consider the protein size, it successfully identified fake recurrent mutations resulting from large protein sizes. These results provide evidences to support the specificity of our method to recover functional mutations.

### 3.4 Mutation types

Based on mutation types, all the somatic mutation can be divided into missense mutation, nonsense mutation, silent mutation, frame shift Insertion and frame shift Deletion. Taking TP53 as an example, we checked the mutation type distribution between two groups of TP53 mutations: (1) functional TP53 mutation as identified by our method and (2) non-functional TP53 mutations. As shown in Figure 3, most of mutations were missense mutations (~65 %). Statistical test did not support any significance difference between two groups of patients. However, nonsense mutation and Frame shift Insert types had great ratio differences between two groups. 16% of functional TP53 mutations were observed to be nonsense mutations while only 7% of non-functional TP53 mutations. The significance based on Fisher’s exact test was at *p* < 0.03. A similar result was observed with Frame shift Insertion, which was 6% of functional mutations and 1% of non-functional mutations (*p* < 0.12). The enrichment of those types of mutations can be explained by the fact that nonsense mutations and Frame shift Insertions are more more likely to disrupt protein function than missense mutation. We also checked other recurrently mutated genes and similar results were observed. These results suggest that predicted functional mutations are more likely to influence the protein function.

**Figure 3:**
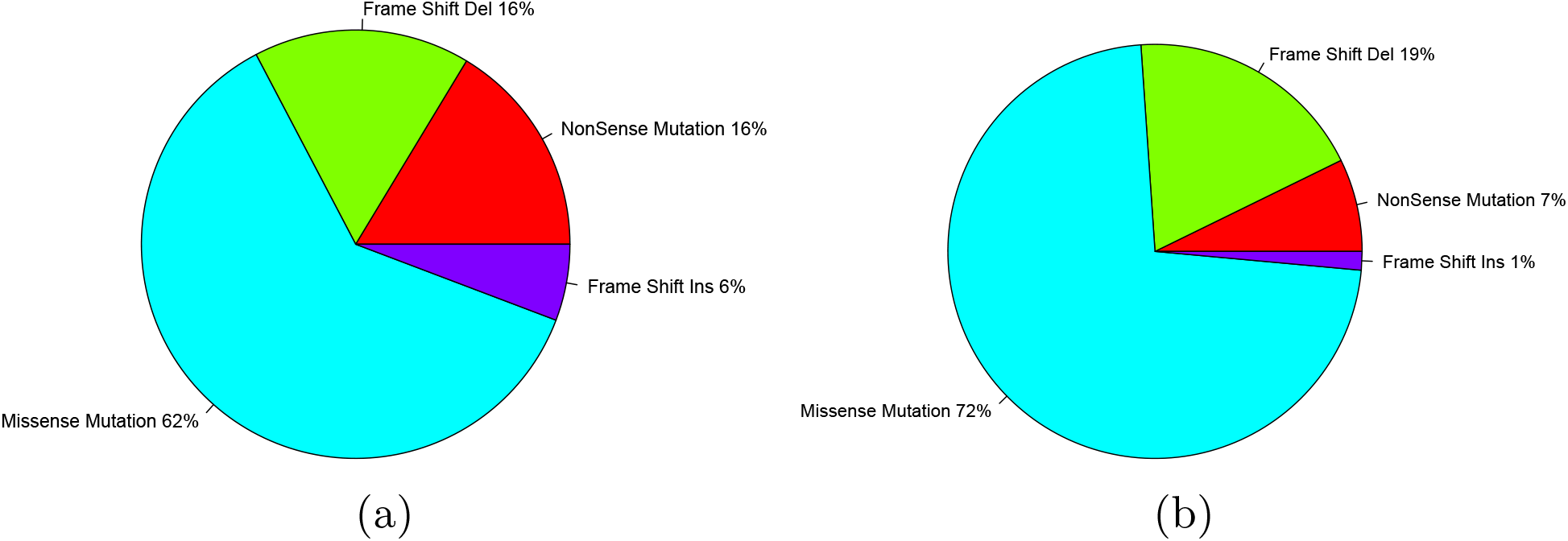
Mutation type differences between functional (a) and non-functional TP53 muations (b)

### 3.5 Cancer-specific gene mutations

Besides of breast cancer, we also applied our method to Ovarian serous cystadenocarcinoma (OV) and Glioblastoma multiforme (GBM) with data from the TCGA project. Figure 4 shows genes with the most recurrent functional mutations of three cancer types. By checking the published literature, we found that most of the predicted genes have been implicated to to have an important roles in oncogenesis [6]. However, we observed quite different mutation preferences among three cancer types. As shown in Figure 4, only TP53 gene is shared by all three cancer types and NF1 gene is shared by OV and GBM. Other genes are preferentially mutated in one cancer type, which can be called cancer-specific mutations. These cancer-specific mutations had higher mutation recurrences and were supposed to take more essential roles in the oncogenesis of a specific cancer type.

**Figure 4:**
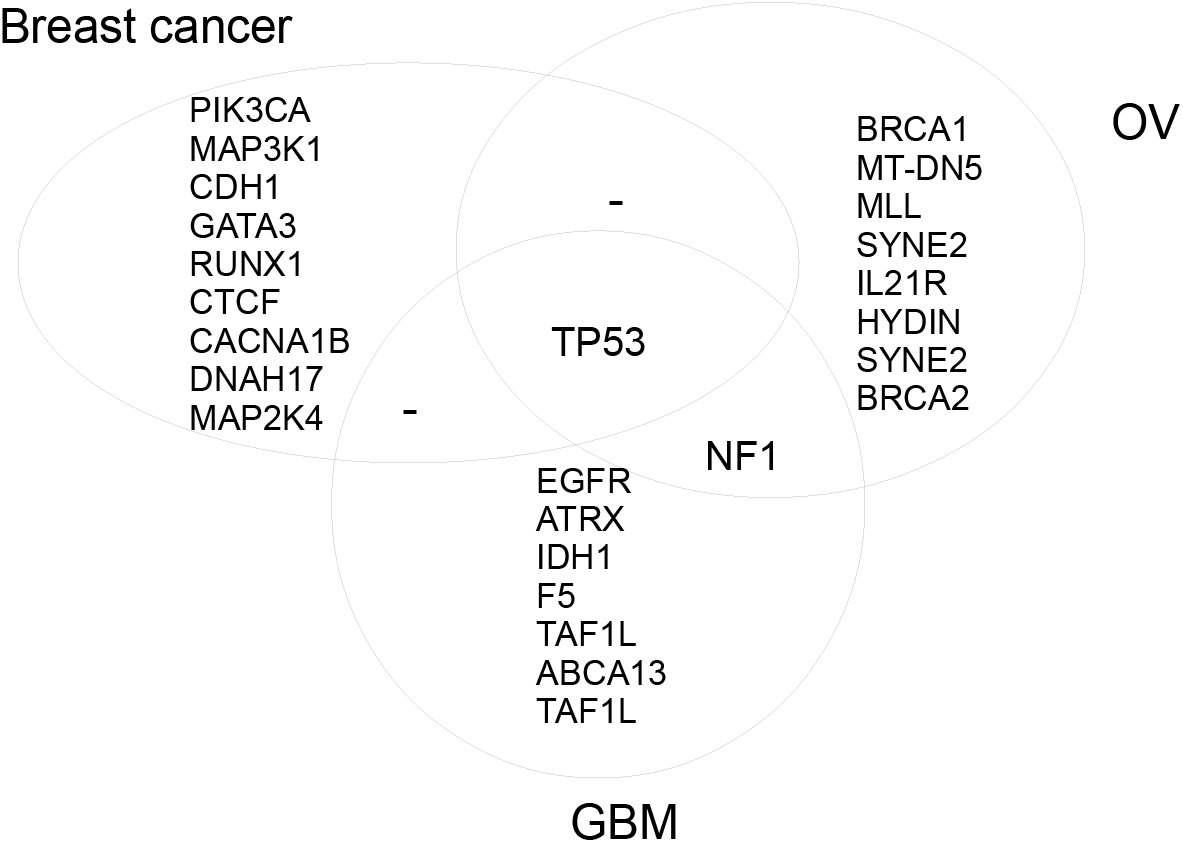
Predicted genes with recurrent functional mutation for three cancers. OV: Ovarian serous cys-tadenocarcinoma; GBM: Glioblastoma multiforme

Cancer-specific mutations are not necessary to indicate non-function in other cancers. One example is BRCA1 gene. BRCA1 was predicted to be mutated in 17 out of 441 patients for OV, ranking as the third most recurrently mutated gene. Based on our definition, BRCA1 is supposed to be OV-specific. However, we also observed functional mutations to BRCA1 in breast cancer. The roles of BRCA1 in the oncogenesis of breast cancer have been widely reported [30] and researches even suggest that patients with BRCA1 mutations have a possibility of 50%-80% to develop breast cancer before the age of 70 [31]. However, mutations to BRCA1 is not supposed to be the most important causal reasons due to the relatively low recurrence, which was at 6 out of 516 patients. BRCA1 is proposed to be involve in many cancer associated biological process, including DNA repair, histone ubiquitination and checkpoint control [32, 33]. One possible explanation for its cancer-specific mutation is that the functional defect of BRCA1 is more likely to be compensated in breast cancer.

### 3.6 Enriched functional terms

As described in Section 2, we can perform functional annotations for each of studied patients by functional enrichment analysis to mutated genes of the neighboring patients. At a cutoff of *p* < 0.01, we observed some terms to be associated with large proportion of breast cancer patients (see Figure 5). One example was the term “cell adhesion”, which was associated with 99.4% of patients. Similarly, other extracellular matrix (ECM) related terms such as “extracellular structure organization” and “cell-matrix adhesion”, were also observed to be associated with 34.1% and 22.1% of patients respectively. These observation are in line with reports about ECM for its critical roles in cancer [34, 35]. This result also suggests ECM to be one potential target to develop anticancer medicine. Above, we reported recurrently mutated genes, such as PIK3CA and MAP3K1. Functional annotation supported their roles by enriched GO terms like “phosphoinositide 3-kinase (PI3K) cascade” (30.6% of the patients) and “MAPKKK cascade” (34.5% of the patients). Considering the fact that PIK3CA and MAP3K1 were ranked as the most recurrently mutated genes, PI3K signalling pathway and MAPKKK cascade pathway were supposed to be the most essential pathways during the oncogenesis of breast cancer. Besides these pathways, we also observed enriched terms such as “JNK cascade” and “cell cycle phase”, which were reported to be related to oncogenesis of cancers [1].

**Figure 5:**
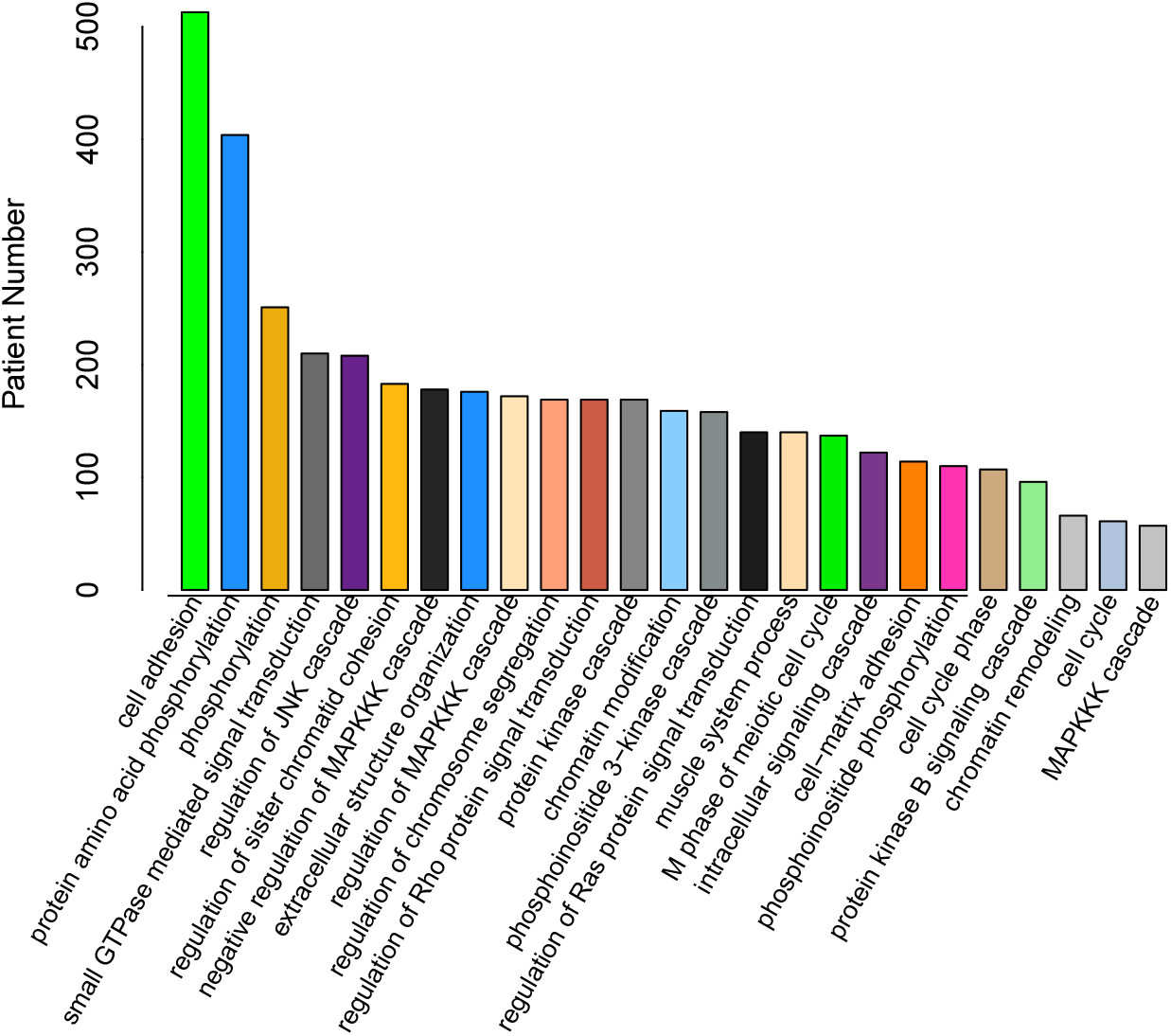
GO terms enriched with breast cancer patients

Based on textbook knowledge and published literature, we could assign some genes to some well-studied pathways or biological processes. For example, TP53 is involved in the p53 mediated DNA damage response [36]; MAP3K1 in the MAPKKK cascade pathway [37]. These terms could be used to evaluate functional differences between patients predicted with or without functional mutations. Taking TP53 mutation of breast cancer as an example, we checked the number of patients with p53 associated term “DNA damage response, signal transduction by p53 class mediator”. Just as showed in Figure 6(a), we observed 6 out of 107 patients with predicted functional TP53 mutation to be annotated with this term while 12 out of 81 non-functional TP53 mutation carriers were annotated with this term. Fisher’s exact test indicated a significant ratio difference at *p* < 0.03. We further checked other TP53 associated terms and observed similar results (results not shown). Besides of breast cancer, we also performed the same evaluation with OV and GBM. Similar results were observed with TP53 mutations in OV. In GBM, we did not observe significant differences, which might be due to the limited number of patients with TP53 mutations, which was only 65 patient. In a way, these results suggest that patients with functional TP53 mutations are less likely to carry other mutations to the members of TP53 associated pathways. In breast cancer, we also checked MAP3K1 mutation with the similar methods for “MAPKKK cascade” term (see Figure 6(b)). Similar to TP53, we observed that the patients with functional MAP3K1 mutations were less likely to carry the mutations to other components of MAPKKK cascade pathways (*p* < 0.01). In GBM, we checked another well-known gene EGFR for that it involved in “epidermal growth factor receptor signalling pathway” (see Figure 6(c)). Similarly, other genes from EGFR signalling pathways were less likely to be mutated in patients with functional EGFR mutations (*p* < 0.08). In summary, all above examples were in line with the popular assumption for gene mutations: mutual exclusivity [13]. However, one contrary example was also observed with predicted PIK3CA mutations in breast cancer (see Figure 6(d)). By checking PIK3CA associated term “phosphoinositide 3-kinase cascade”, more members of PI3K cascade pathway were observed with predicted functional PIK3CA mutation carriers (*p* < 0.02),which was in line with previous report about cooperation of PIK3CA with other oncogenes [38, 39]. This result indicates that mutual exclusivity is not applicable to PIK3CA mutations.

**Figure 6:**
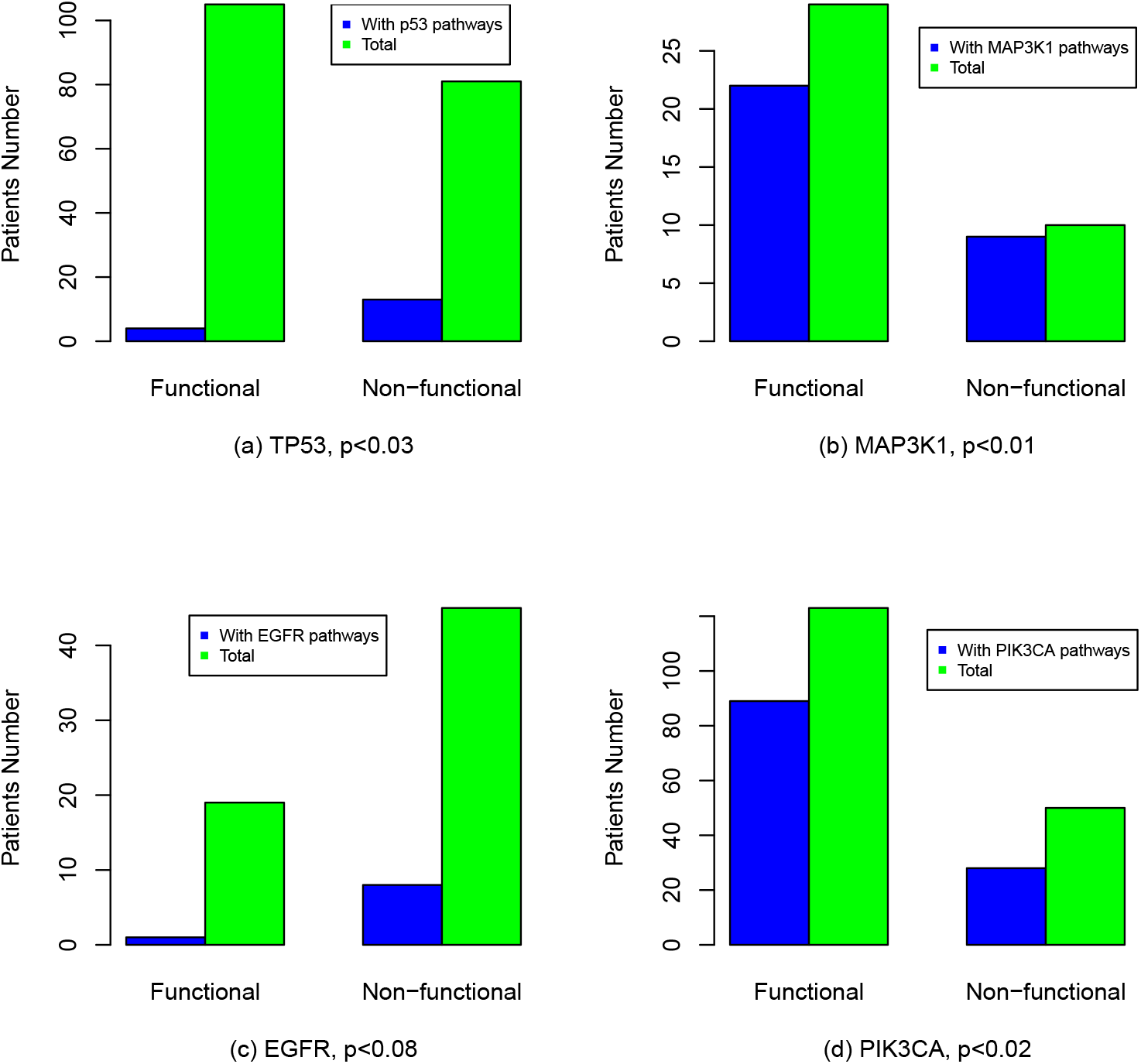
Functional enrichment differences between patients with or without predicted functional mutations.

### 3.7 Mutation network

Based on predicted functional mutations, we also computed the synergy of genes for breast cancer (see Methods). We connected the genes with functional mutations in shared patients and display those genes in a network as showed in Figure 7. Each node is one gene, with node size to indicate its mutation recurrences. We also display the synergy strength with color edges and dark red indicated stronger synergy. In this network, we observe genes, such as TP53, PIK3CA1, MAP3K1 and CDH1, to take hub roles by connecting to other gene. By checking number of connection, we find different genes to take unequal synergy with other genes. For example, 149 genes are connected to TP53 while only seven genes connect to PIK3CA and five genes connect to MAPK3K1. This observation suggests that TP53 can easily have synergy with many genes. This also may be one reason for the observation that TP53 mutation is functional in different cancers. For other recurrently mutated genes such as PIK3A, MAP3K1 and CDH1, they mostly have no connection with TP53, indicating their functional independence to TP53. However, interplays are observed among themselves. This is especially true for PIK3CA, which is also the most recurrently mutated genes in breast cancer. PIK3CA has strong synergy with both MAP3K1 (*p* < 9.1*e* – 22) and CDH1 (*p* < 3.7*e* – 15), another two genes with the mutation recurrence.

**Figure 7:**
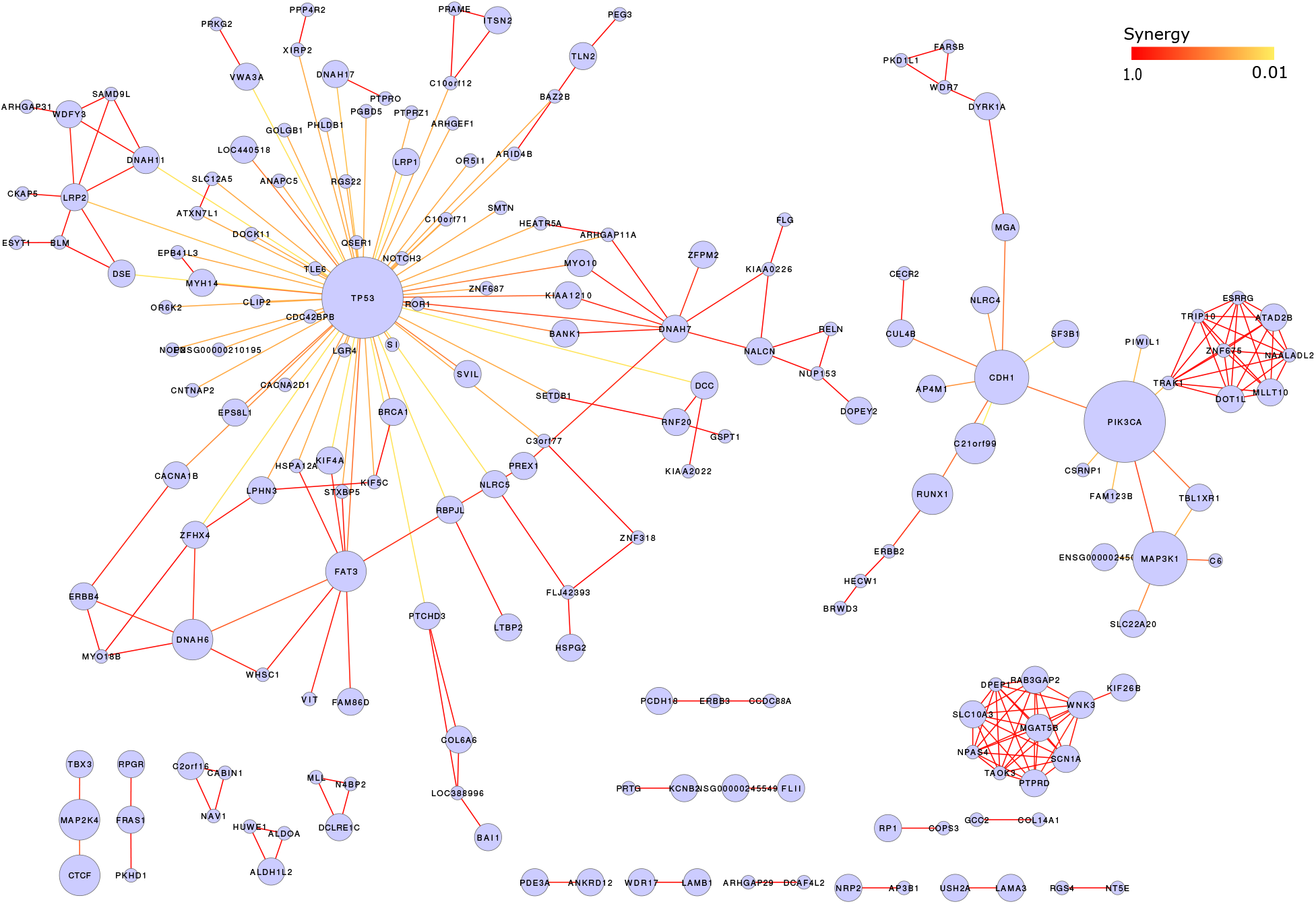
Synergistic network of functional mutated genes

In this network, we also notices some sub-network, whose members were strongly inter-connected with each other but less or no connection to the hub genes, which seems to involve the oncogenesis by their co-mutation during oncogenesis. One example was a sub-cluster including 10 genes, including “WNK3”, “SCN1A”, “PTPRD”, “TAOK3”, “NPAS4”, “SLC10A3”, “DPEP1”, “RAB3GAIP2”, “MGAT5B” and “KIF26B”.

## 4 Discussion

In this work, we propose a computational method for discovery of functional mutation for each studied cancer patient. One basic question is what are functional mutations to cancer. Indeed, different tools have quite different definitions. In the context of this work, functional mutations are mutations that directly or indirectly lead to altered expression of genes on a large scale and that they are the main causal reasons for observed expression patterns of patients. Thus, patients with the same functional mutations are supposed to have similar expression profiles. Following this definition, we transfer identification of functional mutations of one patients to enrichment analysis to mutations in its neighboring patients. This strategy can be called as “guilt by association”, like what we have done in our previous work [40]. By checking the published literature, we can find evidences to support this definition. For example, Langerod A et.al [41, 42] observed characteristic gene expression patterns with patients carrying TP53 mutations. Similar results are available for other cancer genes, such as BRAF in melanoma [24], BRCA1 [43] and PIK3CA in breast cancer [44]. In this work, we also observed preferential distribution of patients with the same mutated genes to sub-clusters by hierarchical clustering (see Figure 1).

The functional mutations are recovered by identifying enriched gene mutations in patients with similar expression profiles. Complexity of mutation profiles make it impossible to cluster all the patients with the same gene mutation into one cluster. Most of the pattern classification methods, such as k-means, support vector machine (SVM), do not consider the sub-structure of samples, which cannot ensure each patient to have the optimal neighbors. With our method, we used a simple methods to find neighbors for every patient one by one, which make it good at recovery of patterns of inner cliques. The utilization of statistical significance also makes this method more robust.

With our prediction, some somatic mutations are not predicted to be functional. It is natural to raise the questions what those non-functional mutations are. One explanation is that non-functional mutations fail to affect the protein function. As an evaluation, we checked protein structure stability between mutated genes with or without predicted function. The folding free energy changes was calculated for each mutation by PoPMuSic2.1 [45]. The energy differences were further evaluated for four examples: TP53, PIK3CA, MAP3K1 and EGFR. However, we only observed significant difference in case of TP53 at *p* < 0.03, which suggested functional mutations to TP53 were more likely to affect its protein structure. For others, no significant differences were observed, which may result from the low accuracy of computational prediction and complexity of mutation to protein function.

For non-functional mutations, another explanation is due to the existence of other stronger functional mutations, which veil the effects of other mutation. It is possible that mutations to one gene may provide cells with proliferation advantage but not enough for cancer, e.g. benign tumours. In this case, other functional mutations are necessary for further oncogenesis and expression patterns of cancer patients will mainly reflect the consequences of latter mutations. Taking TP53 as an example, we observed that 107 breast cancer patients with functional TP53 mutations also carried 17 PIK3CA somatic mutations while 79 patients with non-functional TP53 mutations carried 27 PIK3CA somatic mutations. The significance of ratio differences was at *p* < 0.018. This observation suggests that patients with functional TP53 mutation are less likely to carry PIK3CA mutations. If only considering the predicted functional PIK3CA mutations, no patient with functional TP53 mutations carried functional mutations to PIK3CA while 16 patients with non-functional TP53 mutations also carried functional mutations to PIK3CA. By checking other functional mutations, all 79 patients with non-functional TP53 mutations had at least one functional mutation to other genes. In summary, these results suggest non-functional TP53 mutations may result from existence of other functional mutations.

Certainly, our method also faces the problem of sensitivity if not enough neighboring patients are available. Our method requires a minimum number of neighboring patients for enrichment analysis which is not always available. This is especially true for those mutations with low recurrence. With the progress of cancer genome sequencing projects, the number of patients with mutation information will increase, which will provide a better basis for functional mutation discovery.

Mutual exclusivity is one popular strategy used to recover driver mutations [13]. Based on the functional annotation with GO and KEGG terms, we checked this assumption with some well-known cancer genes. We observed success with mutual exclusivity in the patients with TP53, MAP3K1 and EGFR mutations. However, we also noticed one counter case with PIK3CA, which was involved in the PI3K signalling pathway [46]. Mutations to members of PI3K are more enriched in patients with PIK3CA mutations. This result suggests mutual exclusivity to be a good assumption in driver mutation discovery but also faces the possibility of failure.

